# TargetDB: A target information aggregation tool and tractability predictor

**DOI:** 10.1101/2020.04.21.052878

**Authors:** Stephane P. De Cesco, John B. Davis, Paul E. Brennan

## Abstract

When trying to identify new potential therapeutic targets, access to data and knowledge is increasingly important. In a field where new resources and data sources become available every day, it is crucial to be able to take a step back and look at the wider picture in order to identify potential drug targets. While this task is routinely performed by bespoke researchers, it is often time-consuming and lacks uniformity when one wants to compare multiple targets at the same time. Therefore we developed TargetDB, a tool that aggregates public information available on given target(s) (Links to disease, safety, 3D structures, ligandability, novelty,…) and assembles it in an easy to read output ready for the researcher to analyze. In this manuscript, we will present the methodology used to develop TargetDB as well as test cases.

## INTRODUCTION

With the rising availability of genome-wide association data (GWAS)[1], proteomics[2,3], CRISPR[4–6], RNAi[7], … the list of potential targets for a certain disease grows rapidly. In this context, researchers are spoilt for choices when it comes to picking a target for further investigations, yet the failure rate in Phase 2a trials suggests that researchers are having a hard time at selecting suitable targets against which to pitch drug discovery efforts. To help them in this task, a plethora of excellent publicly available resources exist, such as UniProt [8], DrugBank [9], ChEMBL [10], Open-Targets [11], Therapeutic Target Database (TTD) [12], The Drug Gene Interaction database (DGIdb) [13], Target Central Resource Database (TCRD) [14] and many others [15]. While they all provide valuable information, combining all this information in a single place for further analysis or prioritization of a list of targets can become a daunting task. With each data source specializing in different areas such as protein expression, disease association or pharmacology, it requires the researchers to collate and navigate through endless cross-references in order to paint an accurate portrait of a target. Although resources such as UniProt, Pharos/TCRD and Open-Targets already propose some aggregation of data, we propose with TargetDB to complement them with additional information such as structurally enabled druggability assessment, area-specific scoring for agile prioritization and a tractability prediction model.

## MATERIAL AND METHODS

### TargetDB

TargetDB is distributed as a python package and a pre-built SQLite database. The user can also build the database from scratch using a command-line interface in Linux based systems. Details on the database and on how to install the package are available in the supplementary information and on the GitHub page (https://github.com/sdecesco/targetDB)

### Data sources

Data used in TargetDB comes from a variety of data sources. Some data comes from pre-aggregated/processed data from other databases such as UniProt or TCRD. While others come directly from the source API’s such as Human Protein Atlas for protein expression levels and Open-Targets for disease association. The full list of data sources is available in the Supplementary Information (SI).

### Structural assessment of druggability

Fpocket[16] (version 3) was used in order to probe the potential ligandability of queried targets (https://github.com/Discngine/fpocket). For each target in the database, PDB files were downloaded locally and only the smallest biological assembly with a chain representing the target of interest was kept for further analysis. Fpocket was then used with the default parameters and output files read and incorporated in the targetDB database.

### Tractability model

Data collated in TargetDB is then retrieved and used to generate a series of descriptors that are used for: 1) calculate the area-specific overall score, 2) as input for machine learning algorithm in order to predict the target tractability. The final model uses the random forest algorithm from the python package sci-kit-learn[17]. The building of the model is discussed in the results and detailed procedure and code in the form of a jupyter notebook and training/testing data are available in the GitHub repository.

## RESULTS

Once the program and database are downloaded, TargetDB can be run as a Tkinter graphical interface where different modes can be selected (Figure 1): Single Mode, List Mode and Spider plot mode. For each mode, the target(s) of interest need to be specified. In Single Mode, one file will be generated per gene entered and while nothing prevents the user to use this mode for a large number of targets it is best suited for a handful of genes. For a large number of targets, the List Mode is more appropriate as it will produce a single file with several columns that will allow the user to prioritize targets according to many attributes. In Plot Mode, a graphical spider plot representation of a target landscape will be depicted, representing the amount of knowledge on a target in different areas. This plot is also included in the Single Mode output. An example of each output is available in the supporting information.

**Figure 1:**
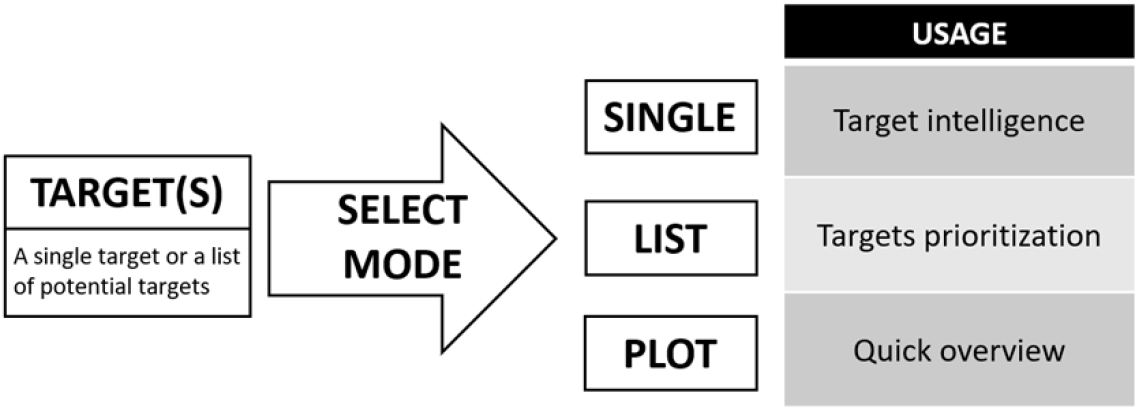
Workflow and usage of TargetDB

**Figure 2:**
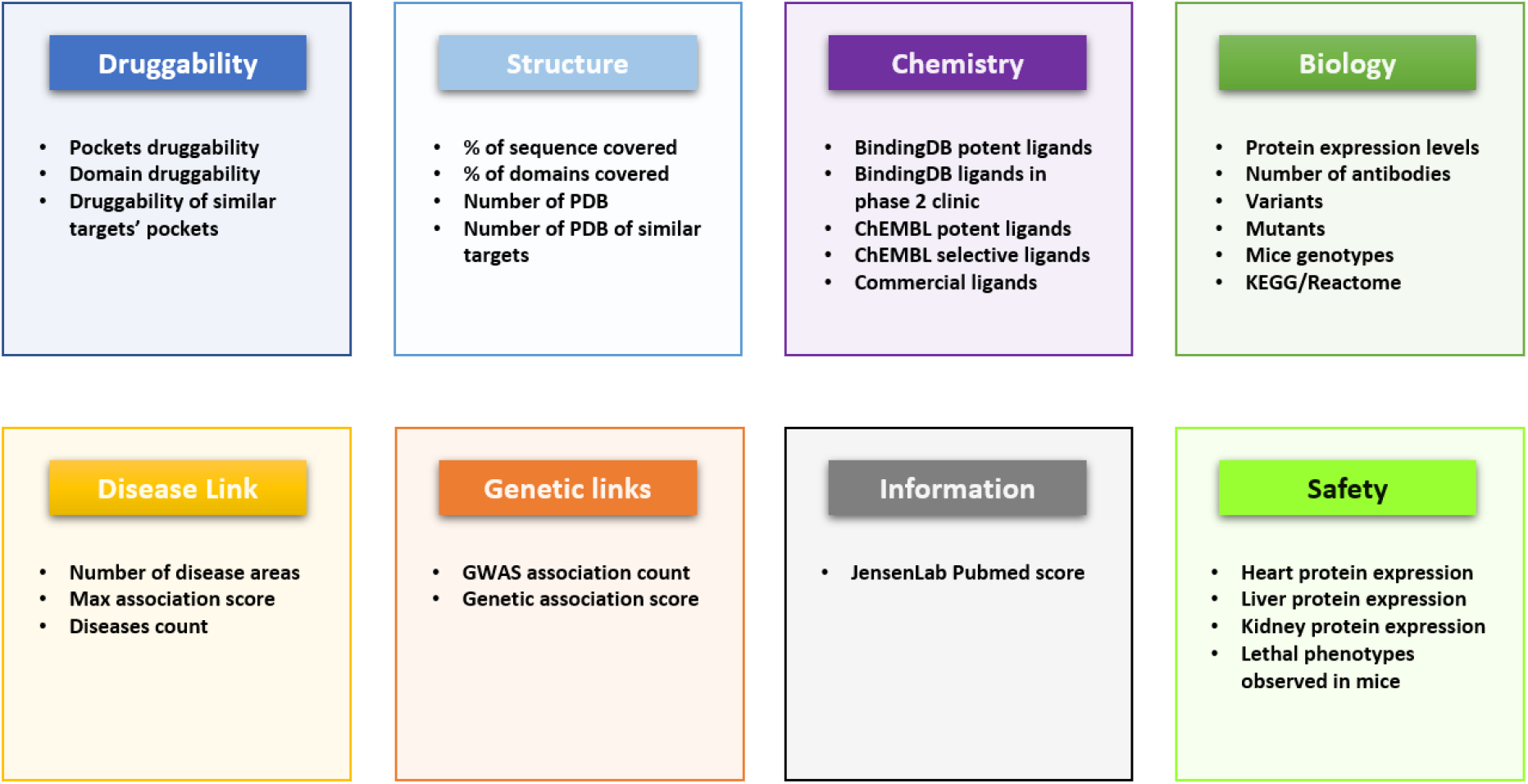
Different features that were selected for the generation of the area-specific scores

### Aggregated information about a specific target

The excel document (see Supporting Information for an example) generated from the database contains several worksheets with different information regarding the target. The main page will contain general information as well as the spider plot. Detailed sheets provided are listed below with a short description.

#### Pubmed search

A pubmed search using the gene name as search term is conducted and the 500 more recent publications are listed in the worksheet.

#### Diseases

This worksheet contains the disease expression (upregulated or downregulated) and GWAS association of the target. This data comes from the Humanmine datasource.

#### OpenTarget Association

Disease associations coming from the Open Targets platform. The individual disease, disease areas and association types scores are displayed.

#### Expression

Protein expression levels coming from the Human Protein Atlas portal. Numerical values can be interpreted as the following: 3=High level of expression; 2=Medium level of expression; 1=Low level of expression; 0=Not observed

#### Genotypes

List of different mouse genotypes for the target of interest with their associated observed phenotypes. Green color identifies genotypes with no abnormal phenotypes observed, while red indicates a genotype with a lethal phenotype observed.

#### Isoforms

List of different isoforms with their associated sequence differences.

#### Variants/Mutants

List of observed variants and mutants along with the sequence change and the effect observed if available.

#### Structure

This worksheet contains a list of all available structures available on the PDB, the code along with the technique, resolution, chain and sequence coverage is listed along with information coming from PDBBind. On top of that, details on domains and their tractability/druggability coming from DrugEbillity is also displayed.

#### Pockets

After analysis of potential binding pocket with fpocket3, the results are imported in TargetDB and are displayed in this sheet. The druggability score comes directly from the fpocket3 algorithm and we refer the reader to the original paper for more details about the method used to generate this score [18]. As a general guideline, a druggable binding pocket would have a score of over 0.5, up to a maximum of 1. If multiple pockets are found for a single structure, a complete list of them will be output. If no druggable pocket is found in the target PDB or no PDB is available for the target, a BLAST search is performed on sequences that have a crystal structure deposited in the PDB. A similar pocket analysis is then performed, and the result displayed in the output document with the identified target as well as the similarity between them.

#### Binding

Bioactivities extracted from ChEMBL contain many different types and while they all provide valuable information it was decided to segregate the data into different sheets: Binding, Doseresponse, Percent Inhibition, ADME and Other bioactivities. The Binding sheet only contains Ki/Kd datapoints. Bioactivities of a given ligand against other targets were collected and used to calculate a selectivity score (Selectivity Entropy – Shannon Entropy[19]), the name of the target for which the ligand has the best bioactivity is also displayed. To provide more information about the ligands, physicochemical properties, as well as the CNS MPO [20] score, are also provided.

#### Dose-response/Percent-Inhibition/ADME/Other bioactivities

Similar to the above mentioned but with different data types.

#### BindingDB/Commercial compounds

Similar to the above mentioned with bindingDB as the datasource. The commercial compounds worksheet contains a link to the chemical suppliers of potential ligand of the target.

### Prioritize a list of candidate targets

The target List Mode report provides the user with more than hundreds of different metrics to define a potential target (a full description, as well as source of these metrics, is available in the SI). To list only a few of these metrics: number of crystal structures in the PDB, ChEMBL bioactive compounds, Open Targets disease associations, number of antibodies, human protein expression levels in tissues,.. Such an abundance of available fields makes it hard to quickly identify a target’s profile or else to pick the relevant parameters for the prioritization process. Therefore, we use a set of rules to define area-specific scores that will help target assessment and prioritization.

Area-specific scores for rapid target assessment. When evaluating potential targets, building an overall picture of a target profile is not an easy task with the information often fragmented in numerous resources. With TargetDB we separated information into eight main categories: Druggability, Structure, Biology, Chemistry, Diseases, Genetics, Information and Safety. Each category is scored from zero to one according to a set of rules (see Supporting Information). Once calculated these scores can be used to generate a spider plot of the target profile to rapidly identify the strengths and weaknesses of a given target. From the few examples in figure 3, it is easy to identify all these targets are well studied and associated with diseases (neurodegeneration), although only some of these have genetic evidence to support the observation and while acetylcholine esterase and beta-secretase 1 are highly druggable and drugged, it is interesting to note that APOE, one of the main risk factors for Alzheimer’s disease [21], does not score well in the druggability and chemistry area, which is consistent with the poor druggability of an apolipoprotein. These well-characterized examples illustrate how this representation allows for a quick interpretation of a target landscape.

**Figure 3:**
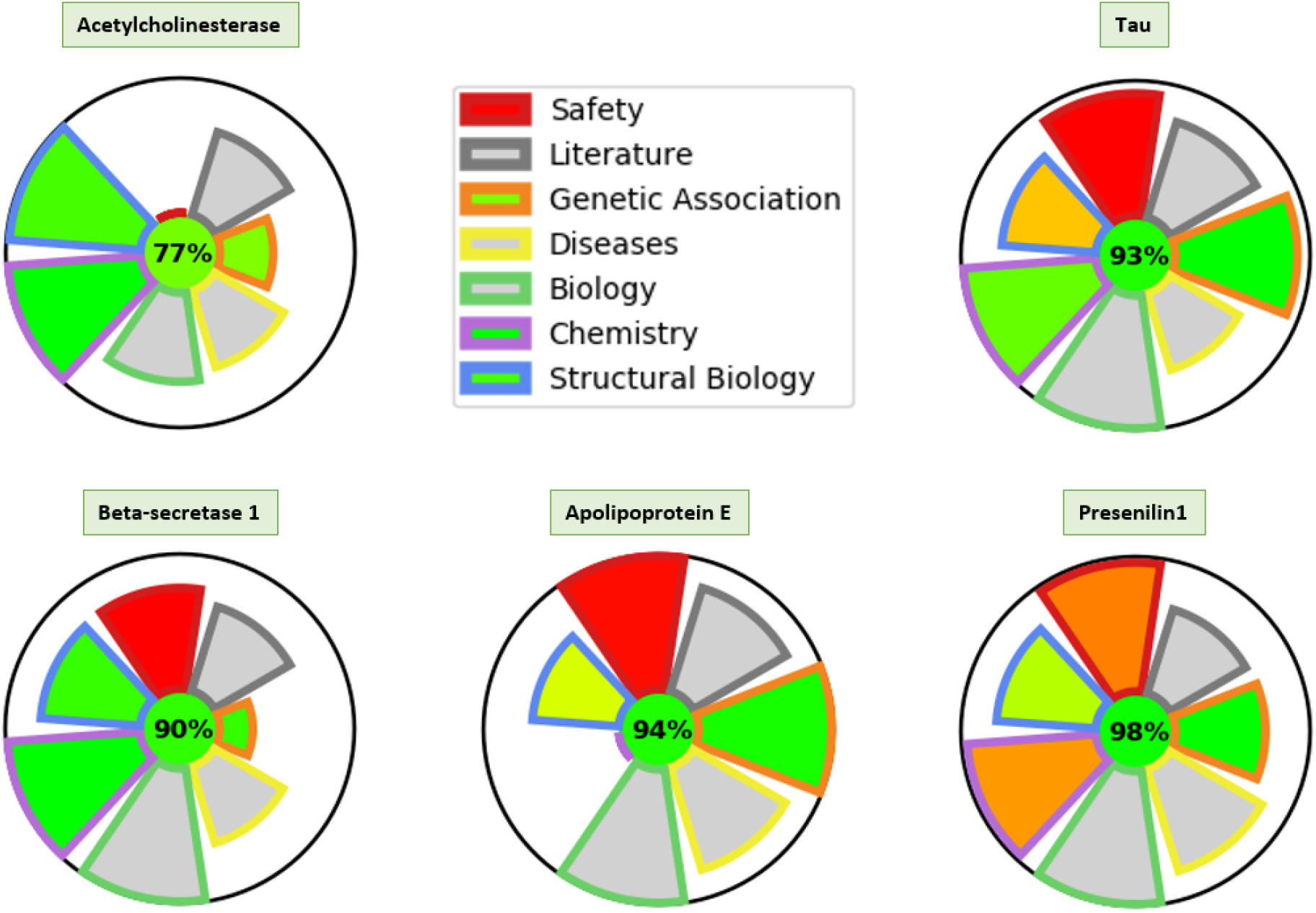
Spider Plot for various targets. The Height of each section represents the amount of information available in that area for this target. The color in the safety, genetic association, chemistry and structural biology are indications of the safety risk, the significance of associations, the quality of the chemical matter and the druggability of potential binding pockets respectively (Green = better quality/safety/… Red = Safety risk/bad quality/…).

#### Multi-Parameter Optimization (MPO) score for target ranking

While ranking targets based on their area score could be used on its own, we also incorporated a customizable MPO score to allow multiple interpretations of the same data. For example, depending on the user interest for a structurally enabled target, it may be advantageous to prioritize targets for which 3D structures are available and with a high druggability score. By simply adjusting the weightings of each category, one can generate a tailored MPO score to facilitate prioritization according to key criteria (Figure 4). Likewise, negative weights can be set to deprioritize high ranking targets and prioritize low ranking targets, this can be useful if, for example, one wants to deprioritize targets with a lot of chemical matter available. The decision on how to set the different weights rely on the user judgment and the specific criteria that are of interest to them. The detailed methodology on how this MPO score is calculated is available in the SI.

**Figure 4:**
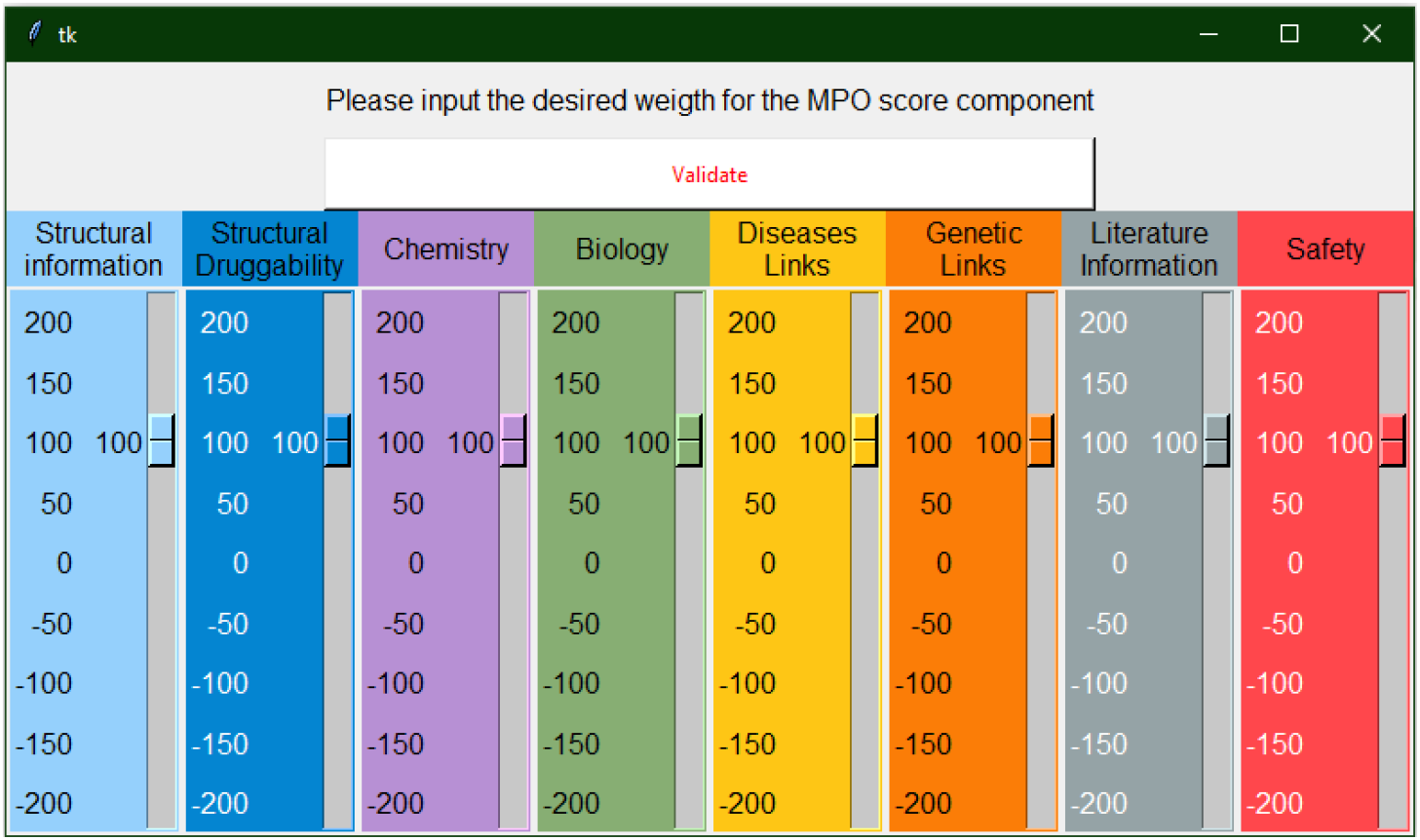
Visual representation of the construction of the MPO Score

#### Target tractability model

To further assist the decision-making process on target tractability, it was decided to evaluate if a model of tractability could be built. With the vast amount of information collected, maybe trends that would allow classification of targets as tractable or not could be uncovered. To do so, several machine learning algorithms were tried, and their performance evaluated to predict target tractability. To train and evaluate the models we had to provide the algorithm with an annotated set of targets, while finding a list of tractable targets is relatively easy, identify a list of untractable targets has proven to be more challenging. We used the DGIdb [13] “clinically actionable” annotated genes as our tractable list of targets(n= 399), while for the negative control we simply selected random targets (n=400) from the list of targets present in TargetDB from which were removed the clinically actionable (n=399) and druggable genome (n=6106) list from DGIdb. This set was then split into a training set (n=560) and testing set (n=240), each containing the same ratio of tractable/untractable targets.

After evaluation of multiple algorithms (detailed procedure as well as the Jupyter notebook available in the SI), the Random Forest algorithm was selected for further optimization (Figure 5). This method provides multiple advantages, such as reducing overfitting, the ability to extract information on features contributing to the decision, and providing an estimate of the confidence of the prediction. The underlying concept of this method is simple: the algorithm creates multiple decision trees, for each decision tree it selects a subset of features from the entire set available, all the decisions from-all the trees are then compiled and a classification based on consensus is made for each target. After feature and parameter optimization the model was evaluated against the test set and was able to accurately predict target tractability 85% of the time. A detailed description of the model performance is available in the SI. The output of this model will then provide two (related) readouts: Percentage of threes predicting the target to be tractable and the tractability class of a target (Tractable [>60%] – Challenging [60%-40%] – Intractable [<40%]).

**Figure 5:**
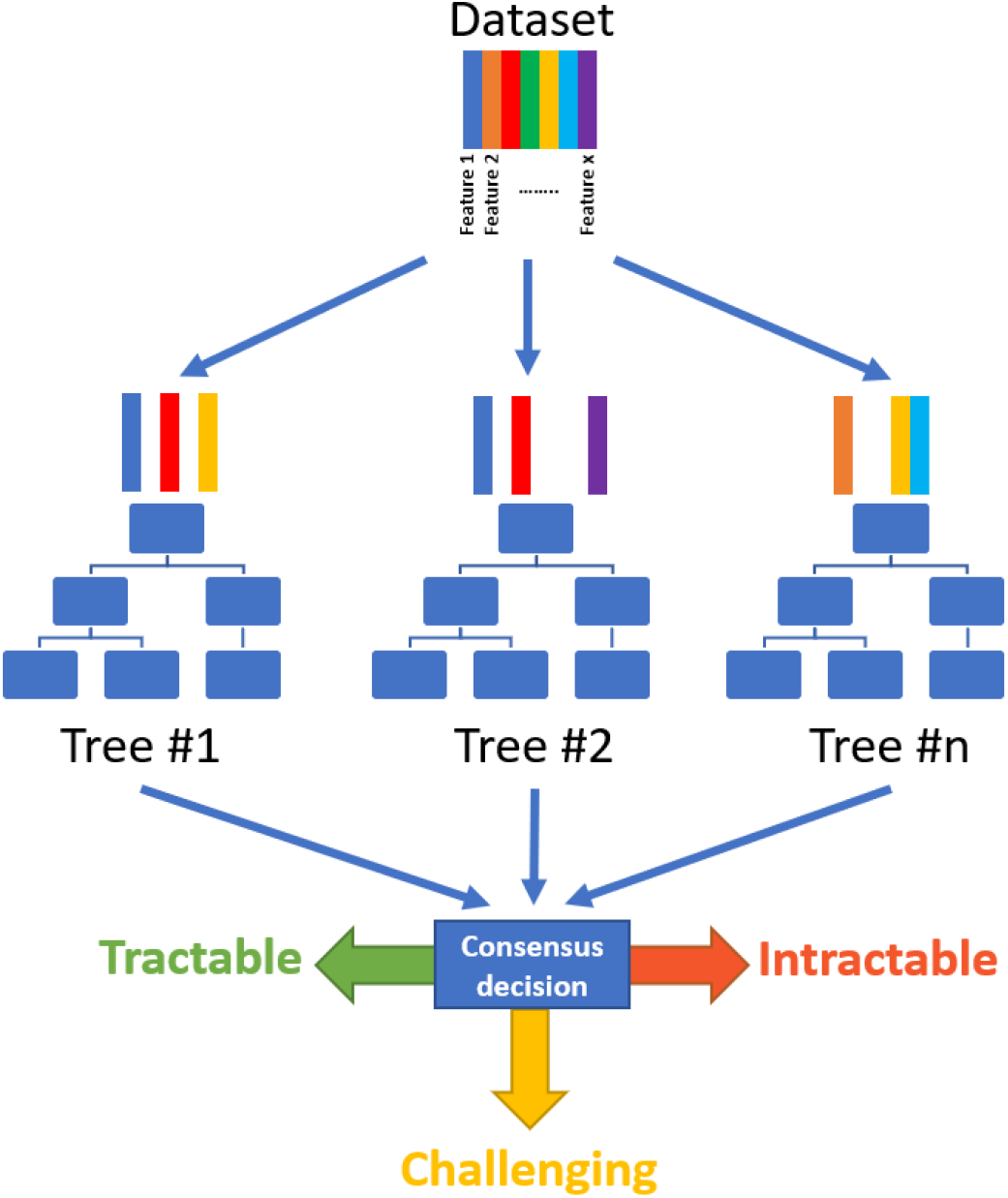
Principle behind the Random Forest Machine learning algorithm

## DISCUSSION

To showcase the application of such a tool, we present here a workflow that was used to prioritize potential targets from a list of genes involved in Alzheimer’s diseases provided by the AMP-AD (https://agora.ampadportal.org). This list consists of 95 targets coming from 6 different teams using computational analysis of genomic, proteomic and/or metabolomic data from human samples. Manual aggregation and collation of information for 95 targets is a time-consuming task, the same results can be achieved in only a few minutes using TargetDB. Once the program started the user has to input the list of targets in the window and select the run mode (single, list, plot), in our case the “List Mode” was selected. Once started, the process will take a few minutes to retrieve all the information in the database and another window will open to allow the user to input each area weight necessary to calculate a custom MPO score. In our case, the following weights were used: Structural information (= 100), Structural Druggability (= 150), Chemistry (= −100), Biology (= 100), Diseases Links (= 100), Genetic Links (= 150), Literature information (= −100), Safety (= 0). The rationale is that we want to select structurally druggable targets, with no or little chemistry available and with strong genetic associations. Biological information and link to diseases are parameters to consider but not essential and we wanted to deprioritize targets with large amounts of literature available. Safety is not considered in the MPO scoring at this time. Once the weights were entered, the program generates an excel spreadsheet with the list of targets and the calculated area-specific, tractability prediction and MPO scores (File available in the supporting information). This spreadsheet can then be used to further refine the selection according to the user’s preferences.

The same list was independently examined by scientists for target selection. 4 targets were selected for further target validation work and early Hit identification. When compared to TargetDB output ranking, 3 of these 4 targets were ranked in the top 10. Assessing these 95 targets took in total a few months and several meetings, it is a good example of how TargetDB could be used to speed up and/or focus the attention on the most promising targets, while not completely discarding the lesser ranked targets for further exploration.

Interestingly, other MPO criteria can be selected depending on the kind of work that is envisioned. For example, a team mainly interested in solving crystal structure might deprioritize targets with a crystal structure solved (Structural information < 0) but still possess druggability potential from close analogs (Structural Druggability ≥ 100) and with some therapeutic rationale (Genetic Links, Disease Links ≥100). These criteria can lead to a ranking massively different than the first one (Table 1).

**Table 1:**
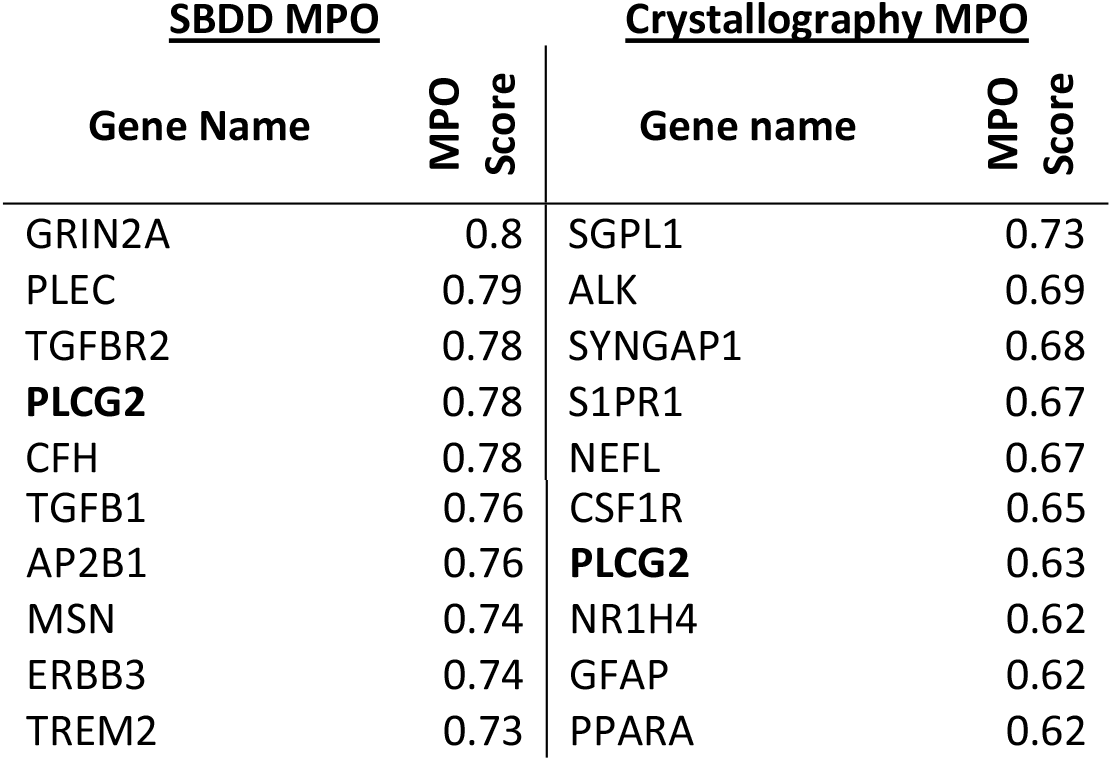
Comparison of the top10 ranked target for two different MPO Score scenario.

| **<u>SBDD MPO</u>** |  | **<u>Crystallography MPO</u>** |  |
| --- | --- | --- | --- |
| **Gene Name** | **MPO Score** | **Gene name** | **MPO Score** |
| GRIN2A | 0.8 | SGPL1 | 0.73 |
| PLEC | 0.79 | ALK | 0.69 |
| TGFBR2 | 0.78 | SYNGAP1 | 0.68 |
| **PLCG2** | 0.78 | S1PR1 | 0.67 |
| CFH | 0.78 | NEFL | 0.67 |
| TGFB1 | 0.76 | CSF1R | 0.65 |
| AP2B1 | 0.76 | **PLCG2** | 0.63 |
| MSN | 0.74 | NR1H4 | 0.62 |
| ERBB3 | 0.74 | GFAP | 0.62 |
| TREM2 | 0.73 | PPARA | 0.62 |

Another application is the prioritization of an entire family of protein. We showcase here how TargetDB was used to rapidly provide an overview of the solute carrier transporters (SLC) family (Figure 6). In less than an hour, we can shortlist potential targets based on their predicted tractability class their MPO score, but also on the potential association with a disease of interest (Alzheimer’s (AD) and Parkinson’s disease (PD) in our case). After further investigations of the top targets, several of them are now under consideration in our institute. This case illustrates how TargetDB can insert itself in the target discovery workflow and expedite as well as standardize the target prioritization process.

**Figure 6:**
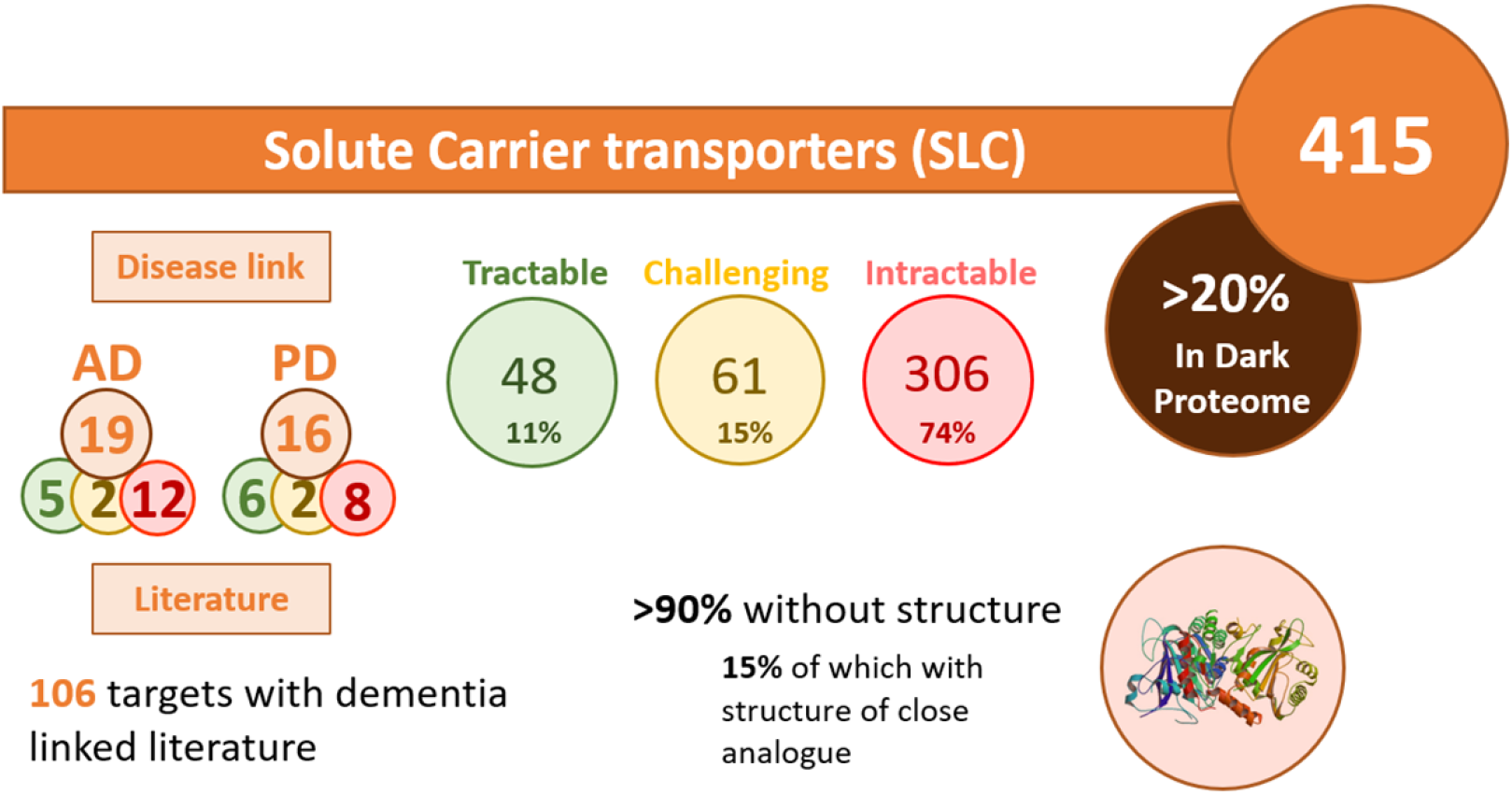
Summary of the analysis of the Solute Carrier Transporters (SLCs) family performed using TargetDB in list mode

## CONCLUSION

In conclusion, we present a tool that allows researcher to extract/combine and standardize outputs from many different publically accessible databases. It allows one to rapidly assess the potential of drug targets. TargetDB is freely available as a python package and detailed installation instructions are available on the project’s GitHub page as well as in the supporting information. While further improvements and additions are already being considered we encourage other users to participate to the project and adding their datasources.

## DATA AVAILABILITY

TargetDB is an open-source collaborative initiative available in the GitHub repository under the GNU GPL-3.0 license (https://github.com/sdecesco/targetDB)

## SUPPORTING INFORMATION

### ACKNOWLEDGMENT

We would like to thank the AMP-AD consortium for providing the target list that was evaluated and the entire Oxford Drug Discovery Institute team for the effort made into evaluating these targets as well as providing useful information for the creation of this tool.

### FUNDING

This work was supported by Alzheimer’s Research UK [registered charity 1077089 and SC042474].

